# MCHM acts as a hydrotrope, altering the balance of metals in yeast

**DOI:** 10.1101/606426

**Authors:** Amaury Pupo, Michael C. Ayers, Zachary N. Sherman, Rachel J. Vance, Jonathan R. Cumming, Jennifer E.G. Gallagher

## Abstract

While drugs and other industrial chemicals are routinely studied to assess risks, many widely-used chemicals have not been thoroughly evaluated. One such chemical, 4-methylcyclohexane methanol (MCHM), is an industrial coal-cleaning chemical that contaminated the drinking-water supply in Charleston, WV, USA in 2014. While a wide range of ailments was reported following the spill, little is known about the molecular effects of MCHM exposure. We used the yeast model to explore the impacts of MCHM on cellular function. Exposure to MCHM dramatically altered the yeast transcriptome and the balance of metals in yeast. Underlying genetic variation in the response to MCHM and transcriptomics and mutant analysis uncovered the role of the metal transporters, Arn2 and Yke4, to MCHM response. Expression of Arn2, involved in iron uptake, was lower in MCHM-tolerant yeast and loss of Arn2 further increased MCHM tolerance. Genetic variation within Yke4, an ER zinc transporter, also mediated response to MCHM and loss of Yke4 decreased MCHM tolerance. The addition of zinc to MCHM-sensitive yeast rescued growth inhibition. *In vitro* assays demonstrated that MCHM acted as a hydrotrope and prevented protein-interactions, while zinc-induced the aggregation of proteins. We hypothesized that MCHM altered the structures of extracellular domains of proteins, and the addition of zinc stabilized the structure to maintain metal homeostasis in yeast exposed to MCHM.

## Background

The potential for significant human exposure to toxic substances is increasing as thousands of chemicals in routine use have had little safety testing (1–3). 4-Methylcyclohexanemethanol (MCHM) is alicyclic primary alcohol used as a cleaning agent in the coal industry. Although health and safety information for this compound is limited, its widespread use in the coal-producing regions of the United States represents a potential hazard to humans and ecosystems. In January 2014, a large quantity of MCHM was spilled into the Elk River in West Virginia, USA and contaminated the drinking water supply of 300,000 people, exposing them to unknown health risks (4). People exposed to MCHM through the contaminated drinking water reported a variety of significant ill effects (5).

MCHM is not easily degraded biologically because of its low reactivity (6). In contrast to other well-studied hydrocarbons, such as cyclohexane and benzene, the effects of MCHM on metabolism are under studied (7). Yeast lines exposed to MCHM exhibited increased expression of proteins associated with the membrane, cell wall, and cell structure functions, while MCHM metabolites mainly induced proteins related to antioxidant activity and oxidoreductase activity (3). With human A549 cells, MCHM mainly induced DNA damage-related biomarkers, which indicates that MCHM is related to genotoxicity due to its DNA damage effect on human cells (3).

Yeast provide an ideal model system to understand the interplay between metabolic pathways involved in the transport, toxicity, and detoxification of MCHM. Further, the use of yeast strains with mutations in various metabolic pathways allows direct evaluation of targeted pathways on the fate and toxicity of MCHM in cells. “Petite yeast”, lines with mutations that disrupt the electron transport chain that produces ATP in the mitochondria, grow more slowly and have smaller cell size than “grande yeast” (wild type). Because these yeast mutants can generate sufficient energy through glycolysis, however, these are not lethal mutations, but provide a slow-growing line to evaluate the metabolic rate and stress response. In addition to their roles in energy transformations, mitochondria are central for the synthesis of amino acids, nucleotides, and heme and Fe-S cluster proteins. Thus, such yeast lines are ideal models to assess the role of mitochondrial function in response to stress.

Petite yeast have different tolerances to chemicals, which may be related to the production of reactive oxygen species (ROS) and mitochondrial function. For example, petite yeast are more tolerant to 4-nitroquinoline 1-oxide (4NQO) than grande yeast when grown on non-fermentative carbon sources that favor respiration (8, 9). 4NQO is metabolized to the active form only in cells with functional mitochondria and petite yeast, without mitochondria and favoring fermentation, are more resistant than wild type yeast. However, petite yeast have higher levels of endogenous ROS and are sensitive to compounds that also generate ROS (10), such as MCHM. Petite yeast are additionally more sensitive to H_2_O_2_ (10, 11) but more resistant to copper (12). Sod1 is the main dismutase that neutralizes ROS in the cytoplasm and in the mitochondria, Sod2, also neutralizes ROS. Thus, the petite yeast-grande yeast pair represent an ideal system to evaluate the role of ROS systems, metal homeostasis, and in MCHM toxicity.

The hydrophobicity of MCHM alters the membrane dynamics, which changes how cells can respond to the environment, including the import and subcellular localization of metals. Metal homeostasis is critical in that metals, such as iron (Fe) and zinc (Zn), play important roles in metabolism as co-factors for enzymes and other proteins, yet, if in excess, induce broad lesions to cell biology through the generation of ROS and by binding to a variety of biomolecules (13). The coordinated activity of metal uptake and sequestration transporters function to maintain metals at optimal levels (Figure 1). For example, there are two Zn transporters located on the cell membrane of yeast. Zrt1 is the high-affinity transporter that transports Zn when extracellular levels are low (14) and Zrt2 the low-affinity transporter (15). Zrt3 transports Zn from storage in the vacuole to the cytoplasm when needed (16) while Zrc1, the Zn/H+ antiporter, and its paralog Cot1 (16, 17) transport Zn into the vacuole from the cytoplasm. Izh1 and its paralog Izh4 are both involved in Zn homeostasis, by altering membrane sterol content or by directly altering cellular Zn levels (18, 19).

**Figure 1.**
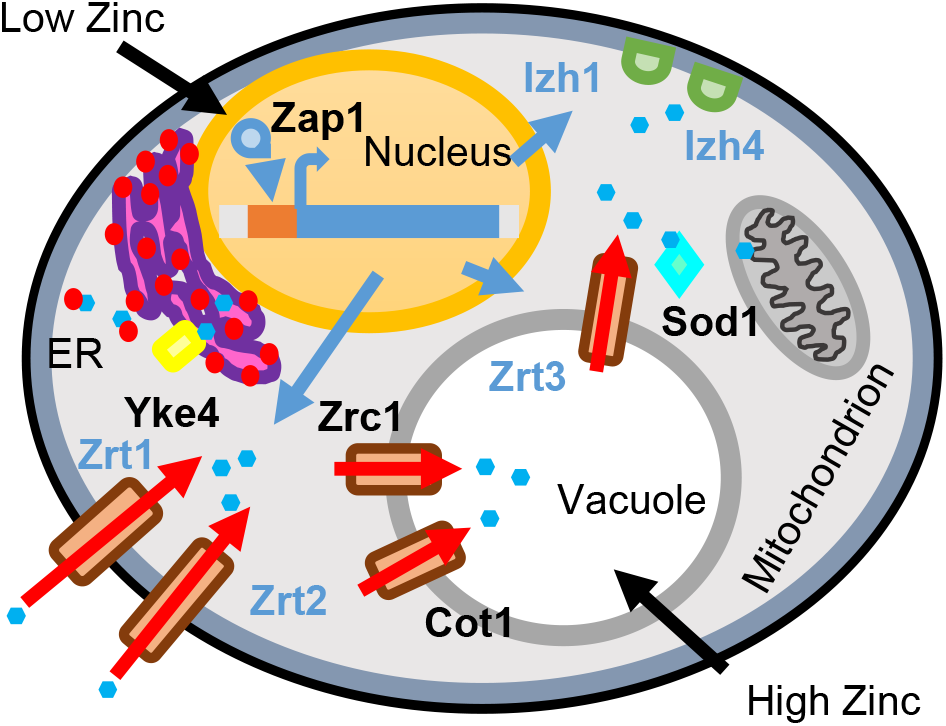
Schematic of yeast cellular response to varying levels of zinc. Under low intracellular zinc levels, Zap1 induces expression of zinc uptake genes. Zrt1, the high-affinity, and Zrt2 the low-affinity transmembrane zinc transporters are localized in the cell membrane. Izh1 and Izh4 localize to the cytoplasmic side of the cell membrane and regulate zinc homeostasis. Excess zinc (blue hexagons) is stored in the vacuole and transported in by Zrc1 and Cot1. Excess zinc stored in the vacuole, and when needed Zrt3 transports zinc to the cytoplasm. In addition to the vacuole, the ribosome (red circles) bind 20% of total cellular zinc (46). Yke4 is the transmembrane transporter at the ER (pink and purple) that transports zinc in both directions. Sod1 is the copper-zinc super oxygen dismutase that is both cytoplasmic and localizes to the inner mitochondrial membrane.

In the current study, we evaluated the impacts of MCHM on petite and grande yeast strains, focusing on metal ion homeostasis and divergent respiratory pathways in these lines as potential mechanisms of MCHM sensitivity. We integrate transcriptomics, ionomics, and GWAS to identify Fe and Zn homeostasis as central to MCHM toxicity and suggest that loss of metal homeostasis underlies ROS damage and MCHM toxicity. While the MCHM has low solubility in aqueous solutions, we propose that MCHM acts as a hydrotrope, altering membrane dynamics and changing how cells responded to nutrients, including import and subcellular localization of metals via transmembrane ion transporters.

## Experimental Procedures

### Yeast strains and media

S96 petite yeast were previously generated from S96 (MATa *lys5*) with a six hours incubation with ethidium bromide. The petite phenotype was validated by failure to grow on glycerol and loss of *COX2* from the mitochondrial genome (9). Yeast knockout strains were previously generated in the BY4741 background (20). The entire coding regions of *YKE4* were knocked out with hygromycin resistance marker in S96 and YJM789 (21). S96-derived strains were grown in minimal media supplemented with lysine, while BY4741 strains grown in minimal media were supplemented with histidine, methionine, uracil, and leucine. S96 and BY4741 are considered in the S288c genetic background while YJM789 is a clinical isolate (22). Crude MCHM provided directly from Eastman Chemical.

### QTL analysis

Isolates of the recombinant haploid collection between S96 and YJM789 (23) were utilized to perform a QTL for MCHM tolerance. A genetic map was constructed and combined with phenotypes collected by growth curves of the segregants in YPD containing MCHM using a TECAN M200 plate reader. Briefly, cultures of segregants were grown overnight then diluted to 0.1 OD_600_ starting concentrations in 200 μl of YPD and either 0 ppm (0 mM) control or 400 ppm (2.8 mM) MCHM. Each segregant was grown in biological triplicate for both control and MCHM treatments in 96-well plates. Both parent strains were grown on every plate to normalize plate-to-plate variation. Plates were grown with constant shaking and OD data was collected every 10 minutes for 24 hours. Differences in control and MCHM saturated concentrations from hours 14-19 of the growth curves were averaged into a single data point to serve as the phenotype for each segregant.

The S96 × YJM789 segregant collection used for genetic analysis contains 126 segregants genotyped at 55,958 SNPs identified by physical location. To perform the QTL analysis as previously described (24, 25), the genetic map was estimated for use in place of the existing physical map. Computational efficiency was also improved by collapsing the 55,958 markers into only 5,076 markers. The R/QTL package based this reduction of markers on all markers that did not recombine and segregate within the population, which it collapsed into individual randomly selected markers within the population. The genetic map was created using R version 3.4.3 and the QTL package version 1.41.6. A custom modification of the scripts contained in Karl W. Broman’s genetic map construction guide was used to output the map, specifically utilizing known physical locations to validate that markers were ordered correctly and to identify yeast chromosome order. The qtl package was used to run the QTL analysis with the maximum likelihood EM algorithm method parameter selected to calculate LOD scores at all loci. Significance thresholds of alpha = 0.05 were applied using 1000 permutations to determine the significance of LOD scores.

### Serial dilution assays

Yeast were serially diluted onto solid media as previously described (24). MCHM and ZnSO_4_ were added to media, autoclaved and cooled to 65°C, which was then poured. Specific experimental conditions varying the concentrations of MCHM, zinc, and yeast strains are outlined in the results.

### Transcriptomics

The RNA-seq of S96 and S96 petite yeast was carried out on hot phenol extracted RNA (26). The raw reads from sequencing are from 16 samples GSM2915204 through GSM2915219, including normal and petite cells. “GSE108873_mchm_fC1_count_table_clean.txt.gz” is the count data generated via Rsubread. “GSE108873_mchm_fC1_DESeq_c2.tsv.gz” is the differential expression data generated via DESeq2. The accession number is GSE108873 (https://www.ncbi.nlm.nih.gov/geo/query/acc.cgi?acc=GSE108873). GO term analysis was undertaken with clusterProfiler (27), an R package that implements methods to analyze and visualize functional profiles (GO and KEGG) of gene and gene clusters. For this, the ORF names from genes up- or down-regulated in each condition were translated to the correspondent Entrez id using the function bitr and the package org.Sc.sgd.db. The resulting gene clusters were processed with the compareCluster function, in mode enrichGO, using org.Sc.sgd.db as database, with Biological Process ontology, cutoffs of p-value = 0.01 and q value = 0.05, adjusted by “BH” (28), to generate the corresponding GO profiles, which were then simplified with the function simplify. The simplified profiles were represented as dotplots, providing up to 15 most relevant categories.

### Elemental analysis

Yeast were grown to mid-log phase in YPD at which time MCHM was added to a final concentration of 550 ppm (3.9 mM). 1.2×10^8^ yeast cells were harvested following 0, 10, 30, and 90 min exposure to MCHM. Samples were split and washed, one set twice with water and the other wash once with 10 mM EDTA to remove metals adsorbed to the extracellular matrix. Four biological replicates for each sample were frozen in liquid nitrogen and stored at −80°C. Cell pellets were digested in 300 μl of 30% H_2_O_2_ and followed by 700 μl of concentrated nitric acid. Samples were heated to 85°C until clear. Sample volume was brought up to 10 ml with HPLC grade water. Digested samples were analyzed in technical triplicate by inductively coupled plasma emission spectrometry (Agilent 5110 ICP, Agilent, Santa Clara, CA, USA) at the following wavelengths for each metal: Ca 393.366, Fe 238.204, Mg 279.553, Na 589.592, P 213.618, and Zn 206.200. Concentrations of elements in digests (mg l^-1^) were normalized to protein levels as determined by Bradford assay.

### Hydrotrope assay

Hydrotrope assays were carried out as previously described (29), with the following modifications. Eggs were purchased and used within one day. Egg whites were separated and diluted 1:6 in 50 mM Tris-HCl pH 7.4. In glass tubes, 3 ml of diluted egg whites were mixed with different concentrations of ATP, MCHM, and zinc sulfate. Samples in the glass tubes were read at 450 nm after 45-60 seconds of incubation at 60°C water bath. All treatments were done in 3-4 replicates and averaged together with the standard deviation shown. Statistical differences were determined using a student t-test.

### Spheroplasting

Spheroplasts of the BY4741 yeast strain were prepared as described (30) with the following modifications. HEPES was used as the buffer and 100 mg 20T zymolase was added per OD unit (OD_600_ multiplied by the volume of culture); spheroplasts were incubated for 1 hour at 30°C. In a 96-well plate, 5 replicates of each treatment were recorded: empty well, spheroplast media only, no treatment, 0.1% SDS, 0.5% sterile distilled water, 10 mM sorbitol, 1 mM ATP, 10 mM NaXS, 10 μM ZnSO_4_, 550-1000 ppm (3.9 to 7.1 mM) MCHM. Using a spectrophotometer, the absorbance at 600 nm was recorded every for 15 hours at room temperature.

### Microscopy

Yeast with proteins tagged at the N-terminus with mCherry under the *TEF2* promoter (31) were grown to log phase and then split into 8 different cultures (4 for treated and 4 for untreated). Once in log-phase samples were treated with 550 ppm (3.6 mM) MCHM for 30 minutes followed by a 20-minute incubation time of 25 μM calcofluor white (Biotium catalog number 29067) on a shaker in a dark room. Then 30 μl of each sample onto a microscope slide pretreated with 40 μL of 250 μg/ ml concanavalin A via pipette onto microscope slides and let sit under a hood for 30 min to dry. The coverslip was placed on top and nail polish around the edges to hold coverslip in place. Cells were imaged on Nikon A1R confocal microscope using a FITC Laser and DAPI laser. Quantitative analysis of pixel intensity to measure the change in expression of the proteins of interest after exposure to MCHM was done with ImageJ on 17-20 cells for each condition. The signal was normalized to untreated yeast and statistical differences were determined using a student t-test.

## Results

Petite yeast have different responses to chemicals because the metabolism of the yeast is shifted away from respiration. Indeed, compared to wild-type (grande) yeast, petite yeast were more sensitive to MCHM (Figure 2). Growth inhibition of petite yeast was affected at 125 ppm MCHM while the growth of wild-type yeast was only affected at 500 ppm MCHM. Yeast have higher tolerances to MCHM when grown in minimal media (YM); however, petite yeast grew less on YM in general (Figure 2).

**Figure 2.**
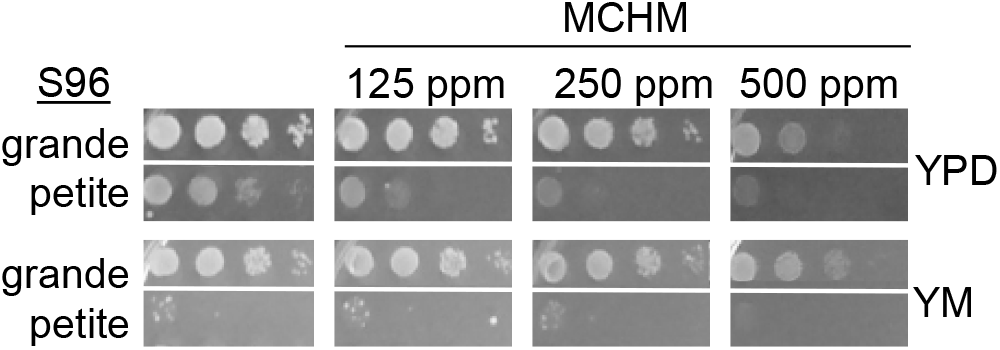
Serial dilution of wild-type S96 and petite yeast in increasing concentration of MCHM in rich (YPD) and minimal (YM) media. Plates were incubated at 30°C for three days and then photographed.

To assess the transcriptional response of petite yeast to MCHM, yeast were grown in both YPD and YM and then treated yeast for 90 minutes with 550 ppm MCHM. 949 genes were differentially regulated across strains and conditions (Supplemental Table 1). Gene expression levels between petite and isogenic grande yeast were similar when grown in YPD media (only six up-regulated genes: cell wall components and iron transporters, and one down-regulated gene: putative mitochondrial protein, Figure 3A), but they were clearly different when grown in YM (131 up and 117 down-regulated genes, Supplemental Figure 1A, Supplemental Table 1). In YM, petite yeast exhibited down-regulated cell wall component genes, and there were also significant changes on small molecule metabolism-related genes (Supp. Figure 1A, 2 and 3, Supplemental. Table 1). MCHM treatment elicited the up-regulation of genes involved in small molecule and sulfur compound biosynthesis, among others, in both petite and grande yeast, while the regulation of other genes differed between strains and depended on the media. For example, genes related to nucleotide and nucleoside metabolism were up-regulated in the petite strain treated with MCHM only when grown in YM (Figure 3, Supplemental Figure 1 and 2, Supplemental Table 1). Genes down-regulated due to MCHM treatment also depended on the media (Figure. 3, Supplemental Figure 1 and 3, Supplemental Table 1). Among the genes with variable expression due to MCHM treatment and/or the use of petite vs. grande yeast were several involved in zinc homeostasis (*COT1, IZH1, IZH2* and *IZH4, ZRT1* and *ZRT2*) and iron homeostasis (*ARN1 ARN2, ENB1, FIT1, FIT2, FIT3, FTR1, GGC1* and *SIT1*) (Figure 3, Supplemental Figure 1 and Supplemental Table 1).

**Figure 3.**
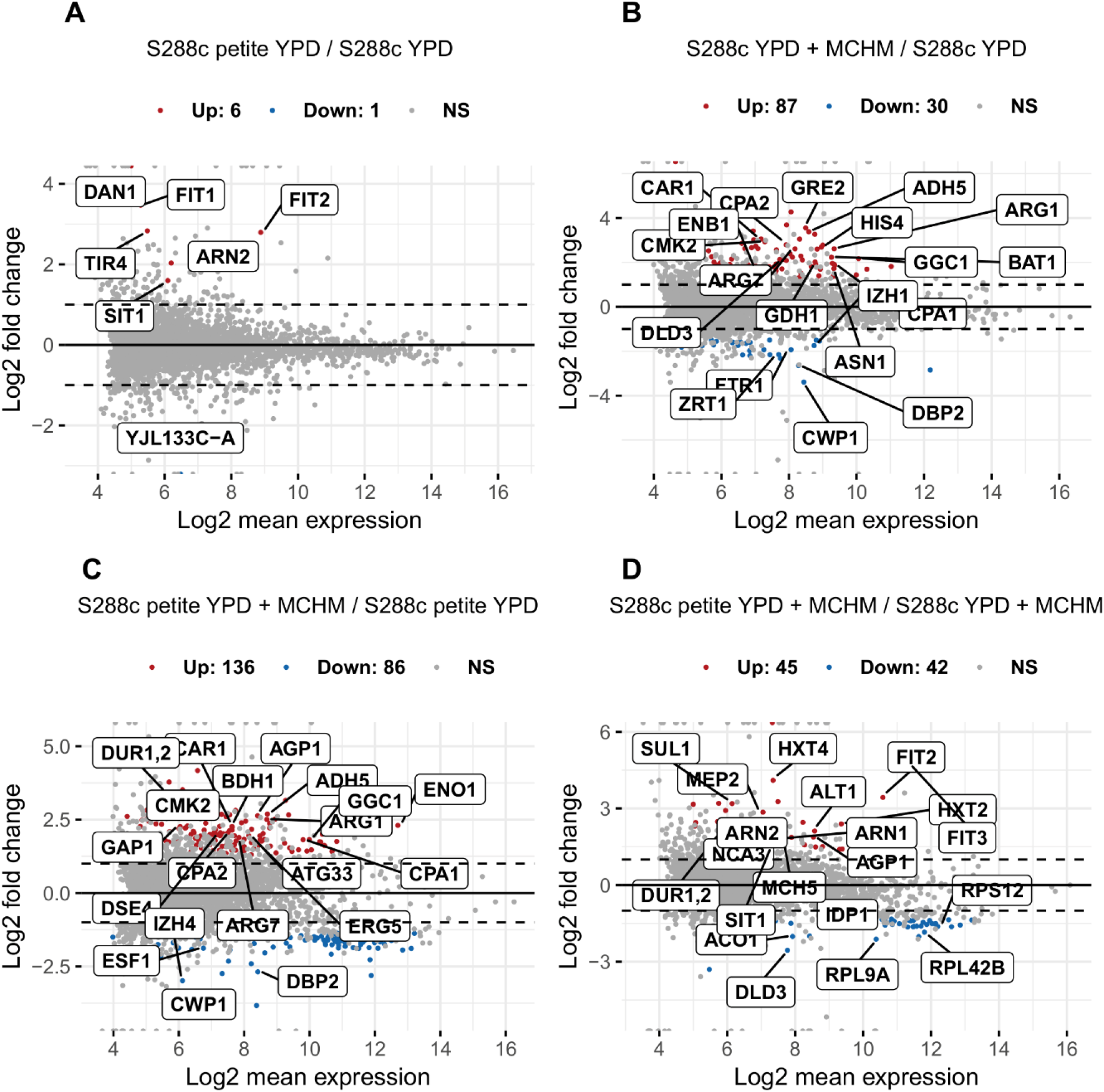
Changes in RNA expression between wild type, petite grown in YPD treated with MCHM. The number of up- and down-regulated genes is noted on the top of each panel. (A) Scatter plots of log fold 2 comparisons of RNA-seq from grande (S96) and petite (S96ρ) yeast grown in YPD. Significantly up-regulated genes are labeled in red and significantly down-regulated genes are labeled in blue. (B) Scatter plots of log fold 2 comparisons of RNA-seq grande yeast grown in YPD and with 550 ppm MCHM. (C) Scatter plots of log fold 2 comparisons of RNA-seq from petite yeast grown in YPD and with MCHM. (D) Scatter plots of log fold 2 comparisons of RNA-seq from S96 and S96ρ yeast grown in YPD and with MCHM.

The increased expression of iron transporters was further explored given that mitochondria, the presence of which differs between the two strains, are the site of iron-sulfur cluster protein biogenesis. Strain ionomic profiles were evaluated for yeast strains grown in YPD. YPD was chosen to minimize the differences in growth between grande and petite yeast. Because there were differences in expression of cell wall genes such as *CWP1*, yeast were also washed in EDTA to determine if increased iron or other metals were associated with the cell wall (water wash) or internalized (EDTA wash). There was no difference in metal levels from yeast washed with water alone or EDTA, indicating that the levels reported ions were absorbed into the cells and were not associated with the cell walls (Supplemental Figure 4 and Supplemental Table 2). Iron levels were three times higher in petite yeast than grande yeast, while zinc was 60% lower in the petite strain (Figure 4A). Other elements were typically lower in grande compared to petite yeast except for calcium (Figure 4A, Supplemental Table 2) as yeast mitochondria do not store calcium (reviewed (32)). Levels of sodium, phosphorus, and magnesium were lower in petite yeast (Figure 4A). Copper was below the limit of detection in this analysis.

**Figure 4.**
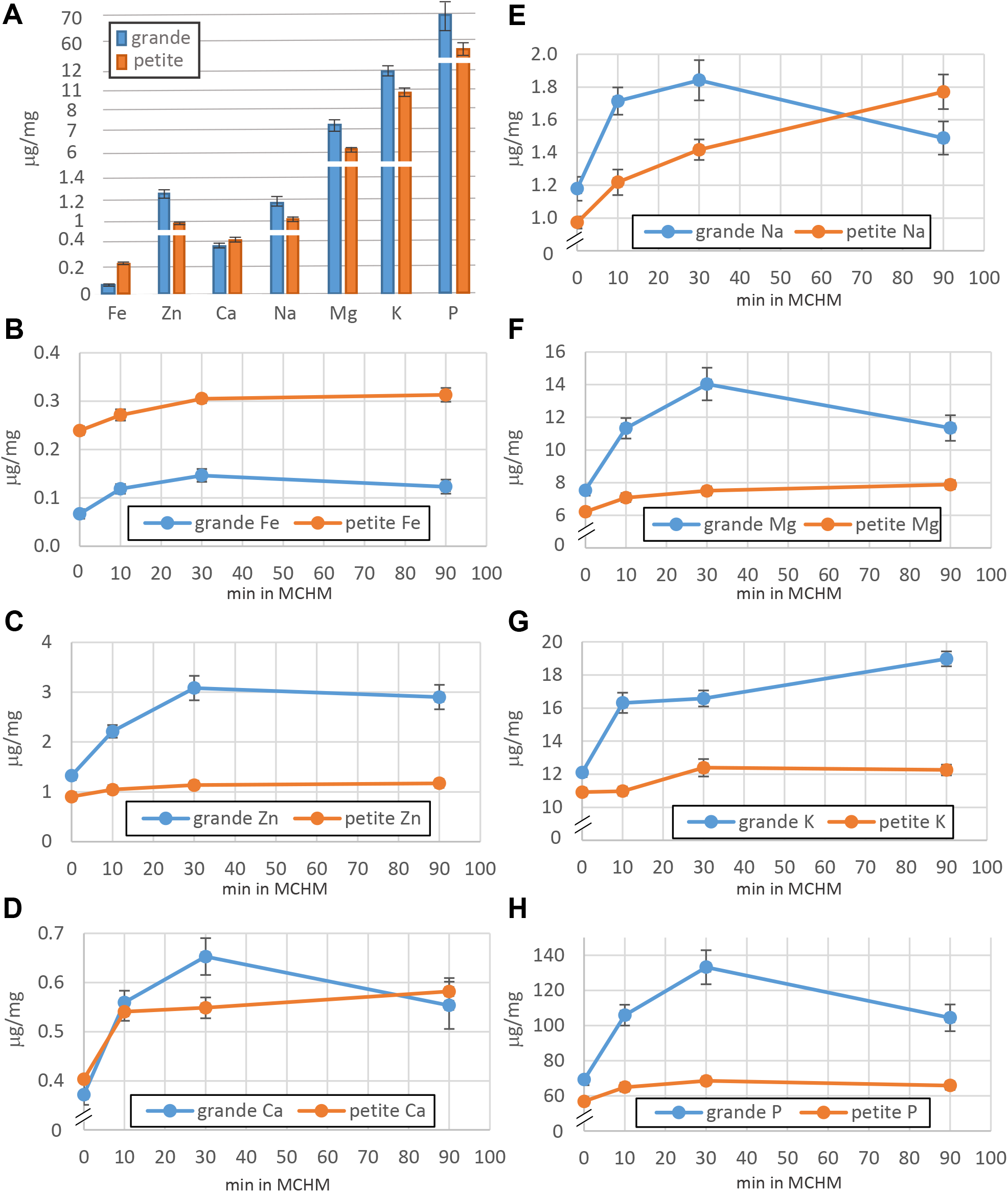
Measurements of metals in yeast treated with MCHM. (A) Levels of Fe, Zn, Ca, Na, Mg, K and K in μg/mg from grande (S96) and petite (S96ρ) yeast grown in YPD. Levels of metals from grande (S96) and petite (S96ρ) yeast grown in YPD with 550 ppm MCHM added for the indicated time. The ions levels measured were (B) Fe, (C) Zn, (D) Ca, (E) Na, (F) Mg, (G) K and (H) F. The standard error is noted on the mean of four biological replicates.

To determine if the levels of metals in yeast were changed when exposed to MCHM, yeast were grown in YPD and then MCHM was added to a final concentration of 550 ppm. The levels of iron did not change for either grande or petite yeast over 90 minutes, although the strain-specific differences noted above, were still notable (Figure 4B). mRNAs encoding siderophores transporters such as Arn1 and Arn2 were expressed at higher levels, as were the Fit mannoproteins, which first bind siderophores in petite yeast in YPD and MCHM compared to wild-type yeast (Supplemental Table 1). In contrast, the levels of zinc increased two-fold in grande yeast but did not change in petite yeast over 90 min (Figure 4C). Calcium and sodium increased with MCHM treatment in both strains with sodium increasing at a slower rate in the petite yeast (Figure 4D and 4E). Potassium, phosphorus, and magnesium also increased in MCHM treatment for grande, but not in petite, yeast (Figure 4F, 4G, and 4H). The levels of these ions are comparable to other studies (33). After 90 minutes of MCHM exposure, five out of the seven ions measured was significantly higher in the grande yeast (Table 1).

**Table 1.** Ionomic profiles (μg/mg protein) of wildtype (S96) and petite yeast strains were grown for 90 min in YPD.

| Element | Wildtype | Petite |
| --- | --- | --- |
| P* | $104.4 \pm 7.5$ | $65.8 \pm 2.2$ |
| K* | $19.0 \pm 0.45$ | $12.3 \pm 0.32$ |
| Ca | $0.554 \pm 0.048$ | $0.582 \pm 0.027$ |
| Mg* | $11.35 \pm 0.79$ | $7.87 \pm 0.25$ |
| Na | $1.64 \pm 0.13$ | $1.77 \pm 0.11$ |
| Fe* | $0.123 \pm 0.014$ | $0.313 \pm 0.014$ |
| Zn* | $2.89 \pm 0.24$ | $1.17 \pm 0.04$ |
| Cu | ld | ld |

\*Denotes significant differences between yeast strains ( $P < 0.05$ ).
ld = level of detection.

There is a significant variation of growth among genetically-diverse yeast strains in response to MCHM. In particular, YJM789, a yeast isolated from a human lung (22), was more sensitive than S96 at 500 ppm MCHM (Figure 5A). Using a segregant collection of YJM789 and S96 that has been used to map genes contributing to differences in phenotypes between strains (8, 23, 24), quantitative trait loci (QTL) analysis was carried out. The growth rate in MCHM of segregant yeast strains was used to assess the association of various parts of the genome with increased growth in MCHM (Figure 5B). Several peaks were noted, but only one broad peak on chromosome nine passed the 95% confidence threshold. Within that peak, we identified *YKE4*, a polymorphic gene that is a ZIP family zinc transporter (34) that plays a role in zinc homeostasis by transporting zinc between the cytoplasm and the secretory pathway (34) and is localized to the ER (31). Yke4^YJM789^ contained two SNPs that change the protein’s amino acid sequence (H5Q and F86L) compared to Yke4^S288c^ (Figure 5C). The H5Q was located in the cytoplasmic signal peptide at the N-terminus while the F86L is at the C-terminal end of the first transmembrane domain using TMHMM (35). To further characterize the role of Yke4, *YKE4* was deleted from S96 and YJM789. Deletion of *YKE4* did not alter growth in the presence of MCHM in these strains (Figure 5D). The zinc levels were measured in these strains. The percentage of zinc was normalized to wild type S96. Both YJM789 and the isogenic *yke4* knockout strains had twice as much zinc compared to S96 (Figure 5E). Yke4 is an ER-localized zinc transporter and plays a role in intracellular trafficking of zinc and did not appear to regulate total zinc levels. There were other genomics peaks in the QTL linked to genetic variation of MCHM response and likely contribute to differences seen between these strains.

**Figure 5.**
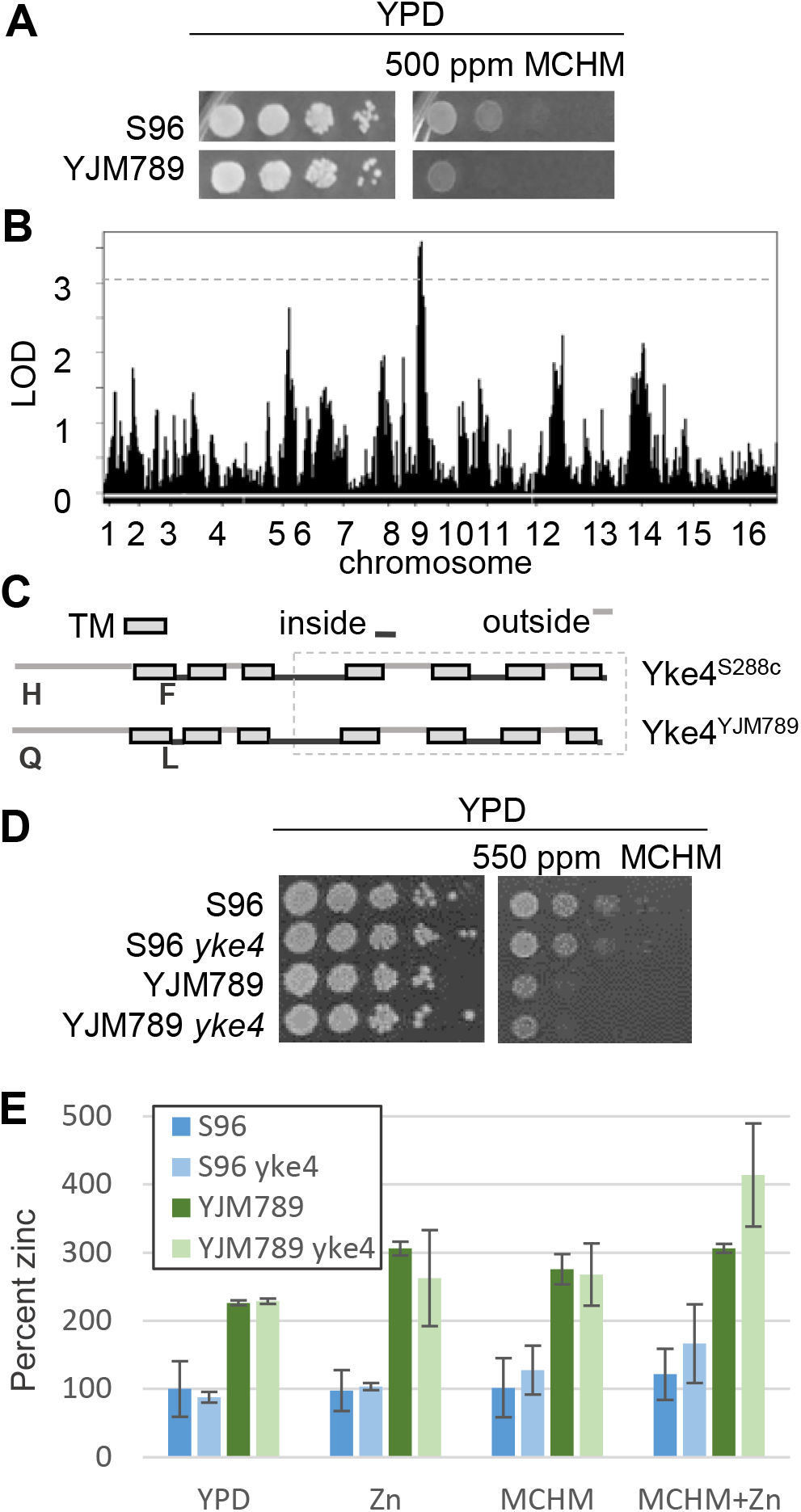
Genetic variation of MCHM response linked to the YKE4 locus. **(A) Serial dilution of genetically diverse yeast on 550 ppm MCHM on YPD**. S96 are from the S288c background and YJM789 is a clinical isolate. (B) Quantitative trait loci analysis of chromosomal regions linked to increased growth of yeast in MCHM from YJM789 and S96 segregants. (C) Diagram of Yke4 describes the transmembrane domains (TM noted with grey box) and inside or outside domains (noted with lower dark grey or upper light grey lines, respectively). Polymorphic residues are noted below in Yke4YJM789 compared to Yke4S288c. The conserved ZIP domain is boxed in a grey dashed line. (D) Serial dilutions of S96 and YJM789 with yke4 mutants on MCHM. (E) Levels of total intracellular zinc from S96, YJM789 and yke4 mutants normalized to S96 grown in YPD and total protein. Yeast were incubated with 5 μM zinc sulfate, 550 ppm MCHM and in combination for 30 minutes before metals were extracted.

To assess the contribution of other proteins involved in metal transport, we utilized the yeast knockout collection to determine the impact of deleting MCHM differentially-regulated genes on growth. This collection is in an S96-related strain background BY4741. In contrast to S96 and YJM789 yeast, the *yke4* mutant in this background was sensitive to MCHM (Figure 6A). There were no significant changes in expression of *YKE4* induced by MCHM (Supplemental Table 1). However, *ARN2* expression was higher in petite yeast and, from ICP-MS analysis, the endogenous levels of iron were also higher. The BY4741 *arn2* knockout was more tolerant to MCHM (Figure 6A). The growth on MCHM of *izh1* and *izh2* knockouts, genes involved in zinc transport that also were differently regulated, was not altered. Iron levels did not change with the addition of MCHM (Figure 4B). However, zinc levels increased in the wild-type grande yeast but not in the petite yeast with MCHM exposure (Figure 4C). We tested whether additional zinc could alleviate the growth inhibition by MCHM. Growth improved with the addition of 10 μM of zinc sulfate in MCHM in both BY4741 and the *yke4* knockout (Figure 6B). However, at higher zinc concentrations (100 μM), all growth was inhibited of all yeast when MCHM was added while zinc sulfate at this concentration alone did not alter growth (Supplemental Figure 5A). Curiously, when zinc was added to YPD without MCHM, the media became slightly opaque. However, after several days, media cleared around yeast colonies. YPD is an undefined media composed of yeast extract, peptone, and agar. Zinc could have induced the precipitation of an unknown compound or compounds that are solubilized by the growth of yeast on solid media; the precipitation of these media components may limit yeast growth at this higher zinc concentration. Therefore, we tested whether yeast knockouts of several known zinc transporters would change the response to MCHM. First, the zinc tolerance of the *zrt1, zrt2, zrt3*, and *zrc1* knockout yeast were tested. Only at the highest levels of zinc sulfate did the *zrc1* mutant grew less than the other strains (Supplemental Figure 5A). Then the addition of 5μM of zinc sulfate completely rescued reduced growth in the presence of MCHM (Supplemental Figure 5B). However, increasing zinc levels to 100μM further suppressed the growth of most of the yeast tested grown on MCHM. The *zrc1* mutant did not grow better at 5 μM and was completely inhibited at 100 μM of zinc in the presence of MCHM. The suppression of growth in higher concentrations of zinc was not due to the toxicity of zinc alone as all yeast tolerated levels up to 500 μM with only *zrc1* mutant displaying reduced growth.

**Figure 6.**
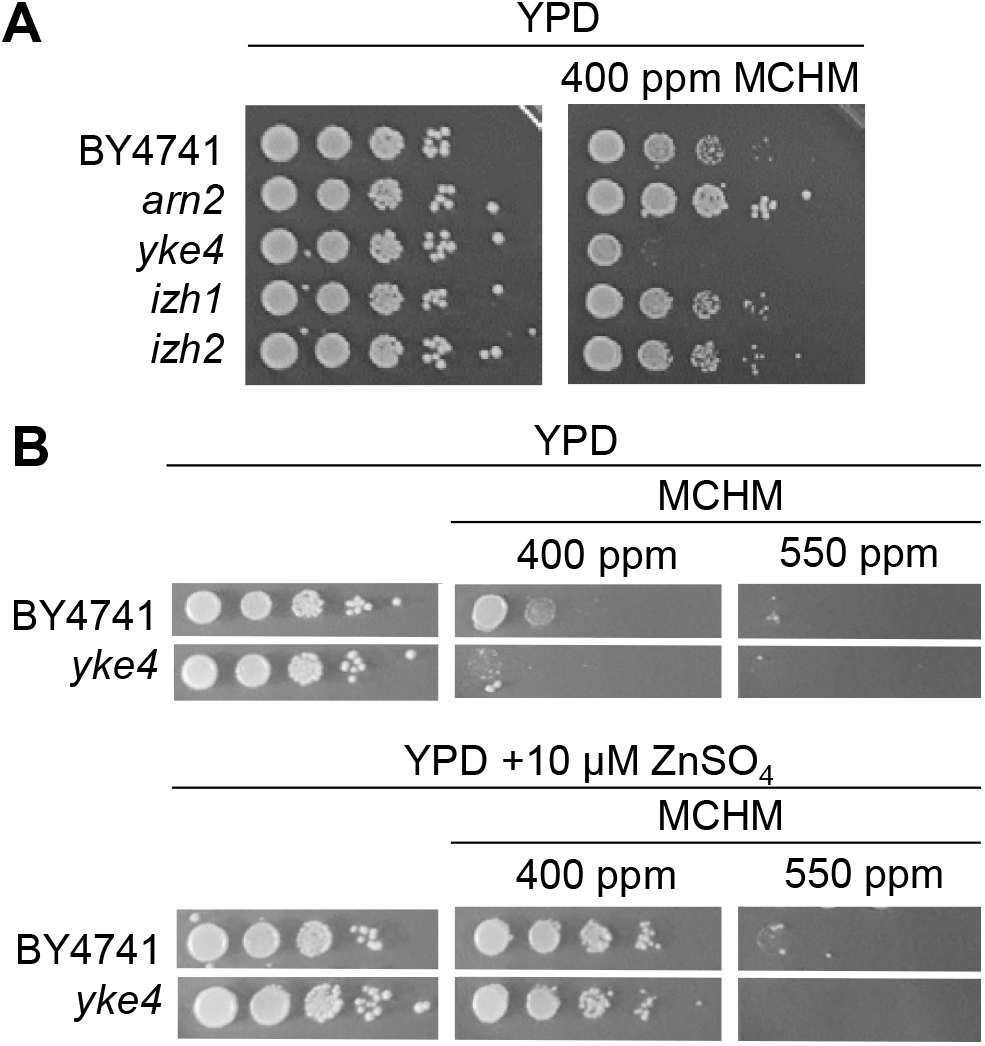
Impact of loss of metal transporters on growth with MCHM and zinc-containing media. (A) Serial dilution of BY4741 and yeast from the knockout collection on YPD with 400 ppm MCHM. (B) Serial dilution of wild type (BY4741) and zinc transporters knocked out yeast grown in 400 or 550 ppm MCHM on YPD with 10 μM zinc sulfate.

MCHM is composed of a saturated hexane ring with a methyl group and a methanol group at opposite carbons (Figure 7A). The methanol and methyl group can be in the *cis* or *trans* conformation. These characteristics of MCHM allow it to act as a hydrotrope, a compound that can solubilize hydrophobic substances in aqueous environments. We thus considered MCHM’s role in altering protein-membrane and protein-protein interactions, which may explain the impacts of MCHM on the transcriptome and ionome. *In vitro* protein aggregation was carried out with sodium xylene sulfate (NaXS), an industrial hydrotrope, ATP, a biologically relevant hydrotrope (29), and MCHM. Compared to no treatment (aggregation was set at 100% for no treatment at 45 seconds), NaXS reduced aggregation to 48% (p= 0.0064), ATP reduced aggregation to 3% (p=0.00025), and 550 ppm MCHM treatment reduced aggregation to 60% (p=0.02). However, at one minute of incubation, MCHM allowed full protein aggregation and was not distinguishable from untreated controls (p=28), while NaXS and ATP continued to prevent protein aggregation (Figure 7B). Levels of zinc sulfate that rescued MCHM-induced growth inhibition increased aggregation by 60% (Figure 7B). Zinc sulfate on its own caused nearly immediate aggregation of protein which was not prevented by the addition of MCHM. We tested if adding MCHM before zinc changed the rate of aggregation. When MCHM was added first followed by zinc sulfate, protein aggregation showed no difference at 45 seconds but was at the highest of all treatments tested at one minute of incubation.

**Figure 7.**
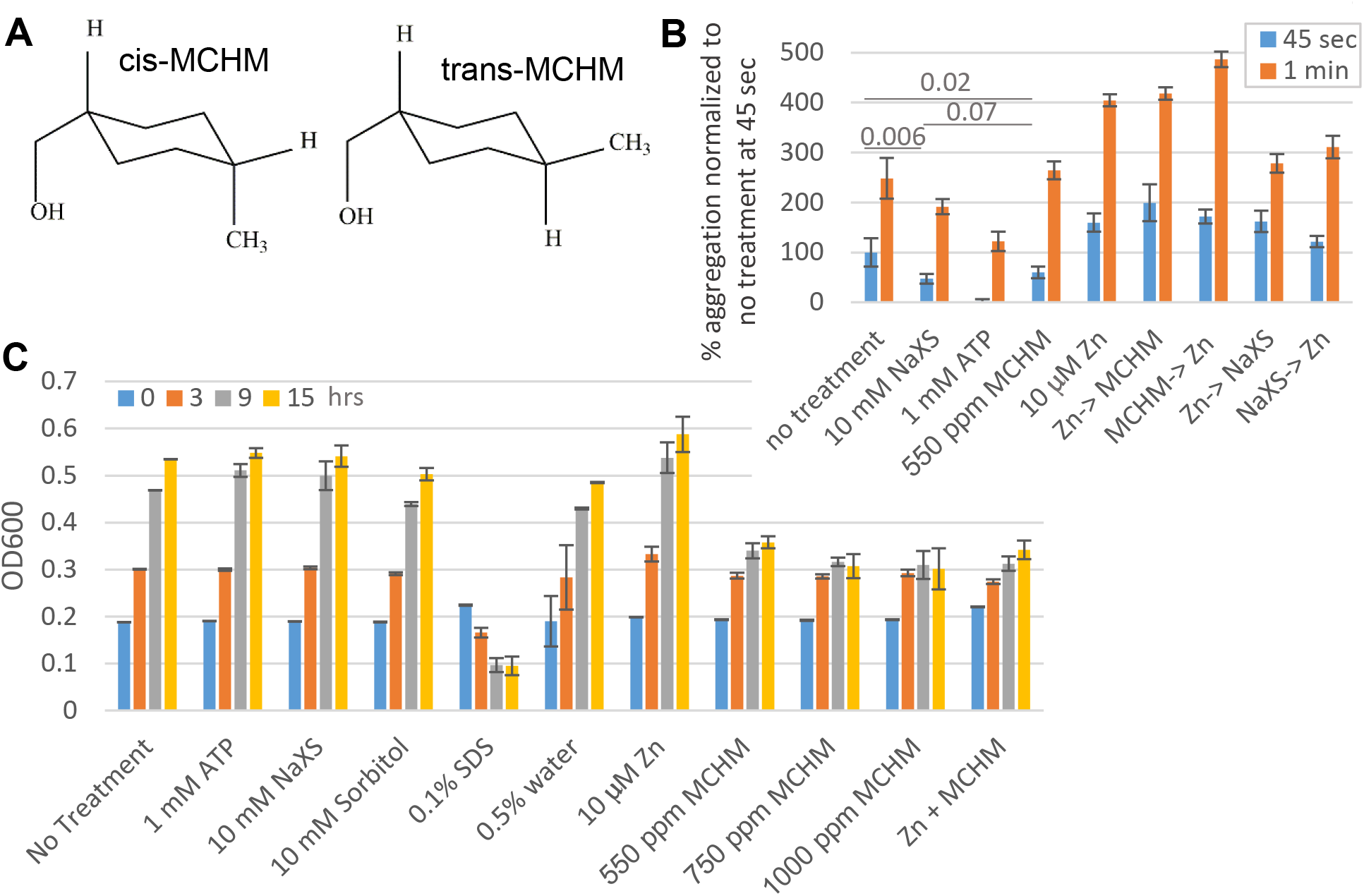
MCHM role in protein aggregation. **(A) Chemical structure of the cis and trans conformation of MCHM**. (B) Timed protein aggregation when exposed to heat. No treatment is compared to 10 mM sodium xylene sulfate, 1mM ATP, and 3.9mM MCHM. The optical density of samples was measured at 450nm after incubation at 60oC. 550 ppm MCHM was added to samples then 10 μM zinc sulfate was added where indicated. (C) Spheroplast yeast were incubated for between 0 and 15 hours with 0.1% SDS, 0.5% water, 10 mM sorbitol, 1 mM ATP, 10 mM NaXS, 10 zinc sulfate and 550-1000 ppm MCHM and samples were read at 600 nm. Biological quadruplicates were averaged and standard error. Relevant p values were calculated using the student t-test.

Yeast have cell walls that protect cells from osmotic stress, and the cell wall can be easily removed to produce spheroplasts. However, unlike plants and fission yeast, spheroplasted yeast continue to divide their nuclei but don’t undergo cytokinesis (36–38) leading to large multinucleated yeast. In this way, we can determine if hydrotropes can cause yeast to lysis when the cell wall is not providing rigid support. SDS, a commonly used detergent, reduced the optical density as yeast cells lysed. Sorbitol provides osmotic support and did not affect yeast growth (Figure 7C). Treatment of spheroplast with known hydrotropes (ATP and NaXS) did not alter the growth of spheroplasted yeast over 15 hours while the growth of spheroplasts treated with MCHM was arrested but did not cause lysis (Figure 7C). The dose-dependent reduction of spheroplasted growth likely mirrors the growth inhibition on plates with MCHM. The growth arrest is reversible as MCHM, as cells will continue growing after MCHM is removed (data not shown). The subcellular localization of Zrt1, Zrt2, Zrt3, and Yke4 was measured as cells were exposed to MCHM. Proteins were tagged at the N-terminus with mCherry (31) and cells were stained with calcofluor white to highlight the cell wall. Fluorescence of each protein remained diffuse and no foci appeared after 90 minutes of exposure (Supplemental Figure 6A). The endogenous promotors were replaced with a common constitutive promoter so the increased expression of Zrt1, Zrt2, and Yke4 were likely at the protein level rather than the mRNA level when exposed to MCHM. Levels of *ZRT1* and *ZRT2* mRNAs were decreased with MCHM (Supplemental Table S1).

## Discussion

The loss of the mitochondrial DNA and treatment with MCHM had pleiotropic effects on yeast. Petite yeast responded to stresses, including MCHM differently than grande yeast. From RNA-seq analysis, iron and zinc transporters were differentially regulated in petite yeast and grande yeast in response to MCHM. In petite yeast, levels of zinc were lower while iron levels were higher. Transcriptomics pointed to metal transporters while genetic analysis uncovered genetic variation in Yke4, an internal zinc transporter as important to MCHM. On non-fermentable carbon sources, *yke4* mutants do not grow in the presence of excess zinc (34), further highlighting that internal zinc transport yeast differed in many other ways. To address how MCHM could affect the wide range of biochemical pathways seen, MCHM was tested and shown to be a hydrotrope *in vitro*, which could exert cell wall or membrane stress.

Petite yeast grew slower and were especially sensitive to the growth inhibition of MCHM. This may be in part due to the altered ionome of petite yeast. This includes higher levels of iron from increased expression of iron transporters and lower endogenous zinc levels. These petite yeast were induced by loss of the mitochondrial genome and petite yeast caused by other types of mutations also had differences in internal metals and regulation in the iron regulome (39). There is an interplay between metal levels, as zinc transporters are also important for responding to high levels of copper (24). Zrt1 protein levels increase in response to high levels of copper (24) and in contrast to the mRNA, Zrt1 under the control of a generic promoter and 5’UTR increased protein expressed by 66% in MCHM treatment. While genetic variation in Zrt2 contributes to copper tolerance (24), Zrt2 protein levels also increased by 26% with MCH exposure. Supplementation with zinc alleviates copper-induced growth inhibition as well as MCHM growth inhibition.

The levels of sodium, calcium, phosphate, and magnesium also increased, suggesting drastic changes at the cell wall and membrane in response to MCHM. Although mRNAs of the iron acquisition pathway were increased in MCHM treatment, there was only a modest change in iron levels. Other stresses, such as starvation induced by rapamycin, also induce the iron regulon (40). Deletion of *arn2* improved growth. Arn2 is localized to the ER and suggesting a role in subcellular localization of iron in the yeast (31). MCHM treated yeast are not starved for iron as *GTL1* and *GDH3* expression was not altered. These Gtl1 and Gdh3 are iron-dependent enzymes that are down-regulated in iron-limiting media and up-regulated in iron-replete media (41). The addition of zinc rescued growth of yeast on MCHM at low levels of zinc (5-10 μM), while levels above 100-500 μM in combination with MCHM drastically reduced growth. Therefore, it appears that the metals have an optimum level to ameliorate the effects of MCHM. Based on RNA-seq and QTL analysis, two transporters of zinc were identified as having an important role in MCHM response. We found no correlation between levels of internal zinc and poor growth on MCHM, possibly because subcellular localization of zinc was critical to growth, rather than absolute levels of the metal. Or there could be a period of adjustment that could not be captured due to the differences timepoints between measuring metal levels and growth. Metal levels were measured at 30 minutes of exposure while growth was measured after two days. *YKE4* expression levels between YJM789 and S96 are not different in unperturbed cells (42) and there are no SNPs in the 5’UTR or 3’UTR (43), suggesting that the polymorphisms in Yke4 itself contributed MCHM sensitivity in addition to other genetic differences. While BY4741 *yke4* was MCHM sensitivity, deletion in S288c and YJM789 had no effect. There are hundreds of genetic differences between BY4741 and S288c and thousands in YJM789 (44, 45). Other yeast strains have the H5Q polymorphism and a subset of those also have the F86L polymorphism. To separate the role of transcription and 5’UTR dependent, the promoter and 5’UTR of *YKE4* was replaced. With MCHM treatment, Yke4 protein levels increased 20%. 582 proteins are predicted to bind protein with 20 proteins binding 90% of total cellular zinc (46) and zinc sparing ensures that essential Zn binding proteins have zinc. As YJM789 expresses less Zrt1 protein normally (24), perhaps the internal levels of zinc and the ability of the yeast to quickly redistribute the zinc in MCHM may alter their ability to grow.

Hydrotropes in cells prevent protein aggregation, but unlike surfactants, work at millimolar concentrations and display low cooperativity. In addition to changes in protein levels, post-translational modifications, and subcellular location, changes in protein conformation also regulate protein function. Protein aggregation is generally thought to inactivate proteins. Protein aggregation includes prions, which increase phenotypic plasticity without changing genetic diversity (47). Intrinsically disordered regions of protein can separate proteins without being membrane-bound which is an important step in RNA granule formation (48). Transmembrane domain proteins such as Zrts, Yke4, and Arn transporters have multiple extracellular and intracellular loops that would be also disordered regions. MCHM slowed the aggregation of proteins in an *in vitro* assay, and with zinc sulfate, that induced protein aggregation appeared to increase the rate of aggregation rather than preventing it. MCHM showed similar hydrotrope activity to NaXS, an industrial hydrotrope, but it was not as potent as ATP. Transcriptomics carried out after 90 minutes, approximately one generation in yeast, detected changes in mRNA encoding metal transporters. Zinc is required for both the syntheses of cell walls and phospholipids (34, 49). Exposure of yeast to MCHM increased intracellular sodium levels yet yeast do not actively accumulate sodium (50), further supporting that MCHM alters the protein structures to either increase the bioavailability of ions or transport across cell membranes. MCHM altered the levels of ions in the cell at the earliest time points. Therefore, we conclude that this is likely through altered protein conformation at the cell membrane of many proteins because of the diverse metals that changed during MCHM exposure. Proteins and organelles are increasingly found in altered conformations to change local concentrations of proteins in (reviewed (51)). Inside the cell, MCHM could alter how molecules interact in liquid droplet changing the function of protein functions and metabolites in the cell.

## Acknowledgments

This work was supported by a grant from the NIH (R15ES026811-01A1) to JEGG and the USDA (1007681R) to JRC and JEGG. WVU Imaging Facilities and the Nikon A1R/N SIM-E are supported by the following grants U54GM104942 and P20GM103434. We are indebted to Wallace Marshall for the insightful suggestions of MCHM as a hydrotrope. Xiaoqing Rong-Mullins assisted in uploading the RNA-seq reads to the public database. Maya Schuldiner from the Weizmann Institute shared the mCherry SWAp-TAG collection. Angela Lee from Stanford University shared the yeast knockout collection. The content is solely the responsibility of the authors and does not necessarily represent the official views of the National Institutes of Health

## Author contributions

JEGG designed the experiments and wrote the manuscript. AP carried out RNA-seq analysis. ZNS measured mCherry tagged proteins, tested protein aggregation and with RJV and MCA carried out yeast growth assays. MCA carried out QTL analysis and constructed strains. JRC analyzed results from ICP and assisted in writing.

## Supporting Information

**Supplemental Table 1**. Comparisons of RNA-seq from S96 and S96ρ yeast treated with 550 ppm MCHM for 90 minutes in YPD and YM supplemented with lysine.

**Supplemental Table 2**. Levels of metals from S96 grande and petite yeast treated with 550 ppm MCHM grown over 90 minutes in YPD. The mean of four biological replicates is shown with standard error. Levels of metals washed with water compared to yeast washed with EDTA.

**Supplemental Figure 1. Scatter plots of log fold 2 comparisons of RNA-seq from grande (S96) and petite (S96ρ) yeast grown in YM supplemented with lysine**. Significantly up-regulated genes are labeled in red and significantly down-regulated genes are labeled in blue. (A) Scatter plots of log fold 2 comparisons of RNA-seq petite and grande yeast grown in YM. (B) Scatter plots of log fold 2 comparisons of RNA-seq grande yeast grown in YM and with 550 ppm MCHM. (C) Scatter plots of log fold 2 comparisons of RNA-seq from petite yeast grown in YM and with MCHM. (D) Scatter plots of log fold 2 comparisons of RNA-seq from grande and petite yeast grown in YM and with MCHM.

**Supplemental Figure 2. GO term analysis of genes up-regulated in all pairwise comparisons with single variable between S96 grande and S96 petite yeast grown in YM and YPD with MCHM added**. The color scale of p adjust values are noted and the size of the circle notes gene ratio. The number of genes in each comparison is noted in parentheses.

**Supplemental Figure 3. GO term analysis of genes down-regulated in all pairwise comparisons with single variable between S96 grande and S96 petite yeast grown in YM and YPD with MCHM added**. The color scale of p adjust values are noted and the size of the circle notes gene ratio. The number of genes in each comparison is noted in parentheses.

**Supplemental Figure 4. Levels of Fe and Zn in grande and petite S96 yeast washed with water and EDTA or with only water before metal extraction**. Mean of four biological replicates are shown with standard error.

**Supplemental Figure 5. Role of zinc transporters in MCHM response with supplemented zinc**. (A) Serial dilution of wild-type (BY4741) and zinc transporter knockout yeast were grown on YPD supplemented with increasing concentrations of zinc sulfate. (B) Serial dilution of wild-type (BY4741) and zinc transporter knockout yeast grown in 400 ppm MCHM on YPD supplemented with increasing concentrations of zinc sulfate.

**Supplemental Figure 6. Levels of mCherry tagged Zrt1, Zrt2, Zrt3, and Yke4 proteins change with MCHM exposure**. Yeast were exposed to 550 ppm of MCHM in YPD for 30 minutes. The cell wall was stained with calcofluor white. B. The mean fluorescence of mCherry normalized to untreated yeast for each protein. Quantification for 17-20 cells shown with standard error.

