## Supplemental Figures 1-6 for "MCHM acts as a hydrotrope, altering the balance of metals in yeast"

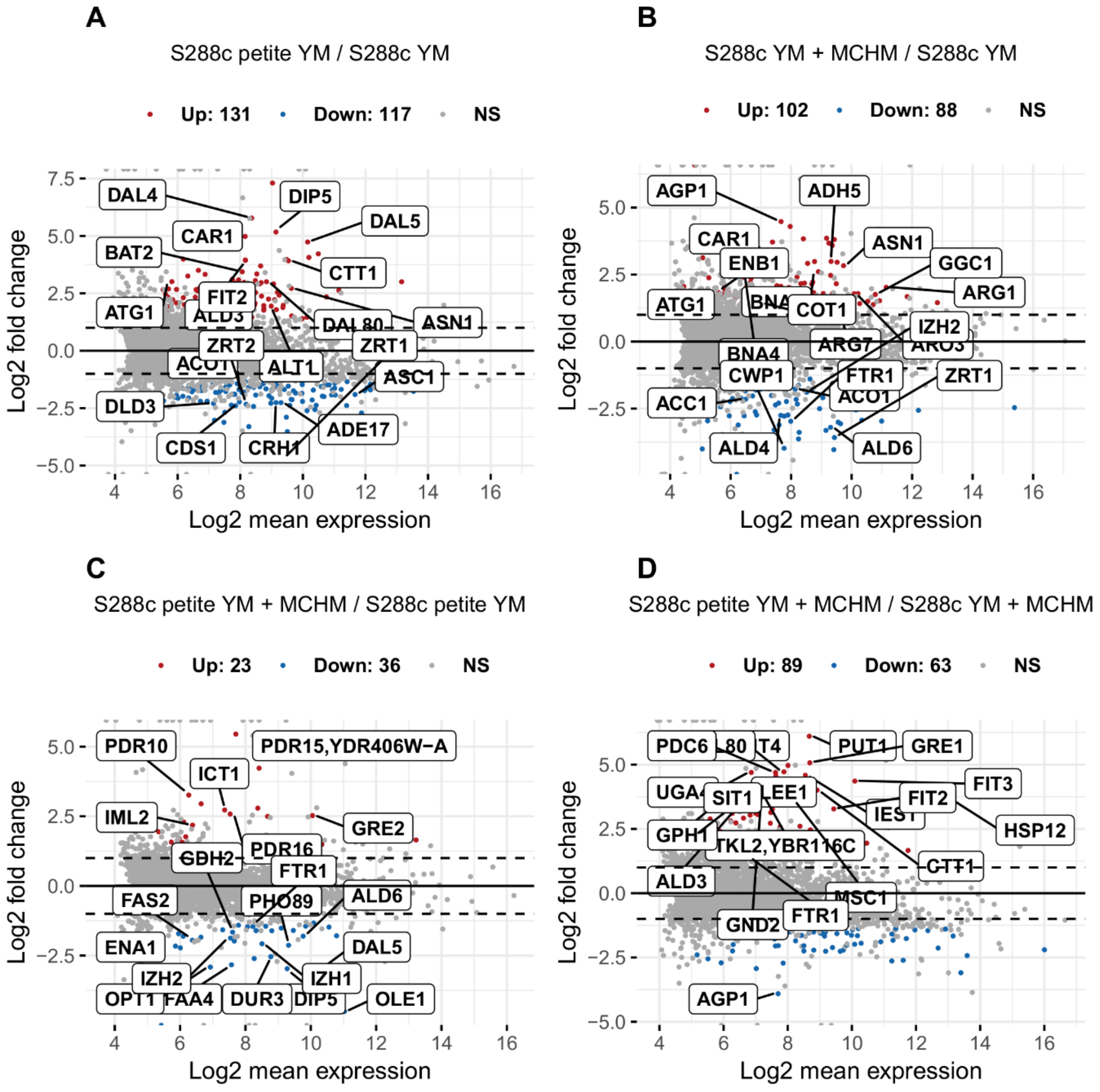


**Supplemental Figure 1 Pupo 2018**

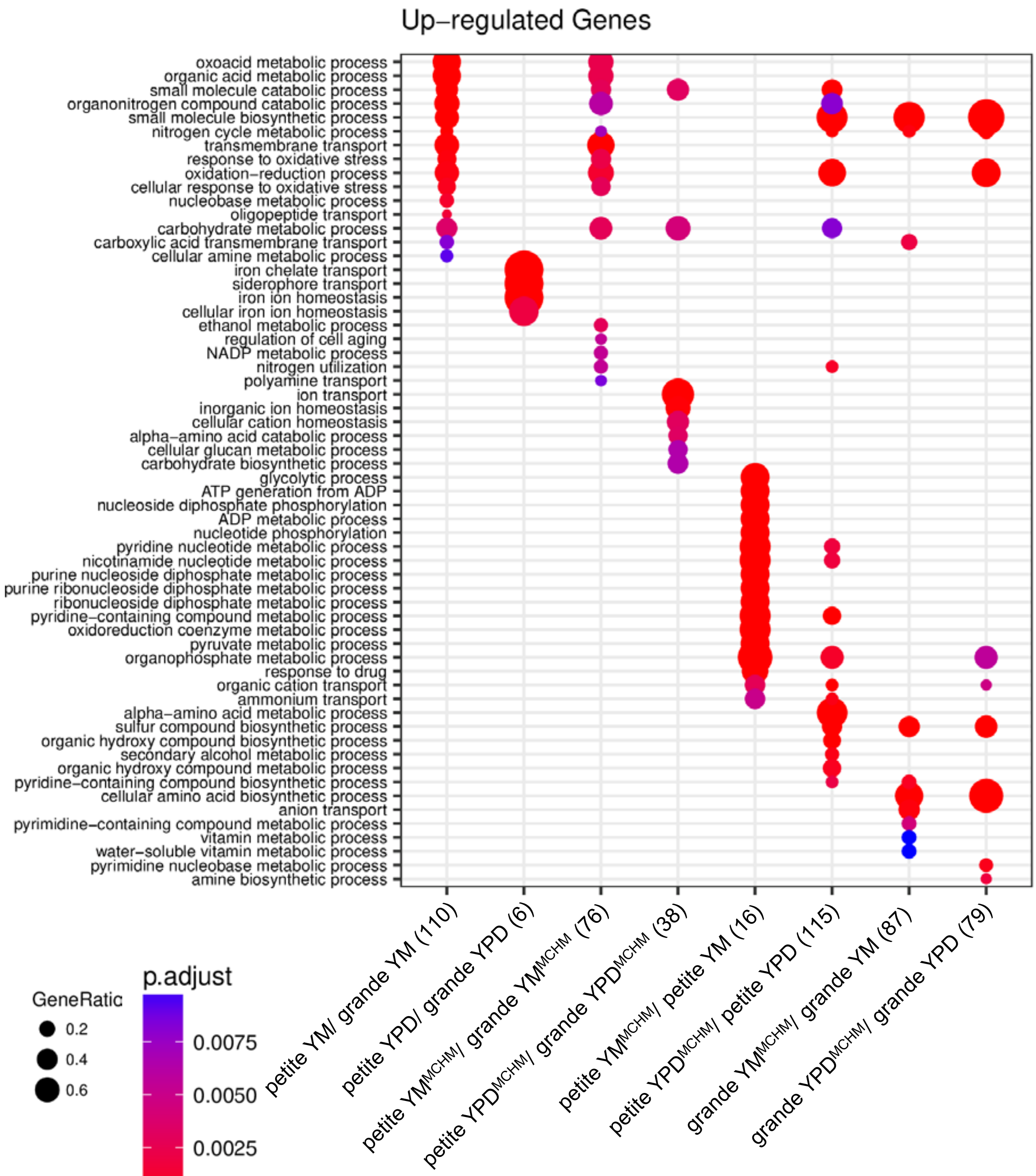


**Supplemental Figure 2 Pupo 2018**

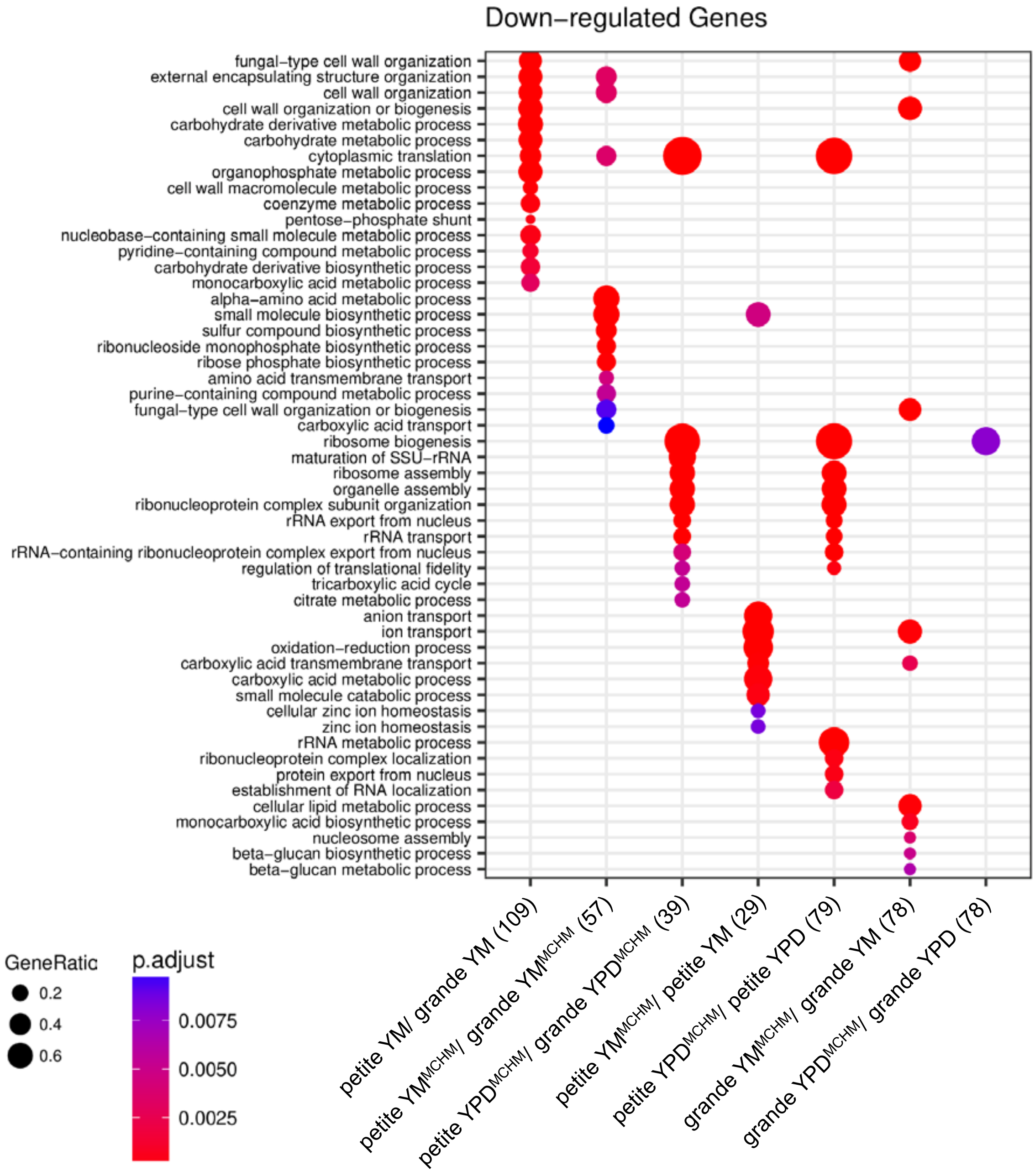


**Supplemental Figure 3 Pupo 2018**

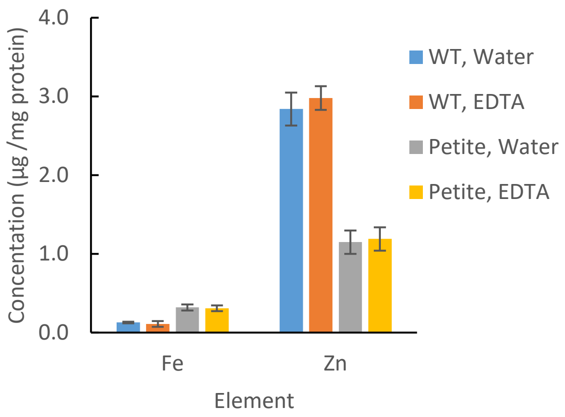


**Supplemental Figure 4 Pupo 2018**

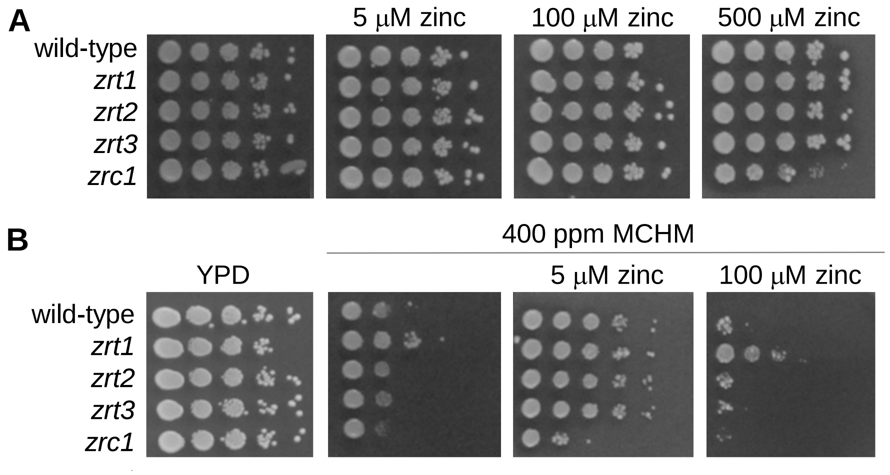


**Supplemental Figure 5 Pupo 2018**

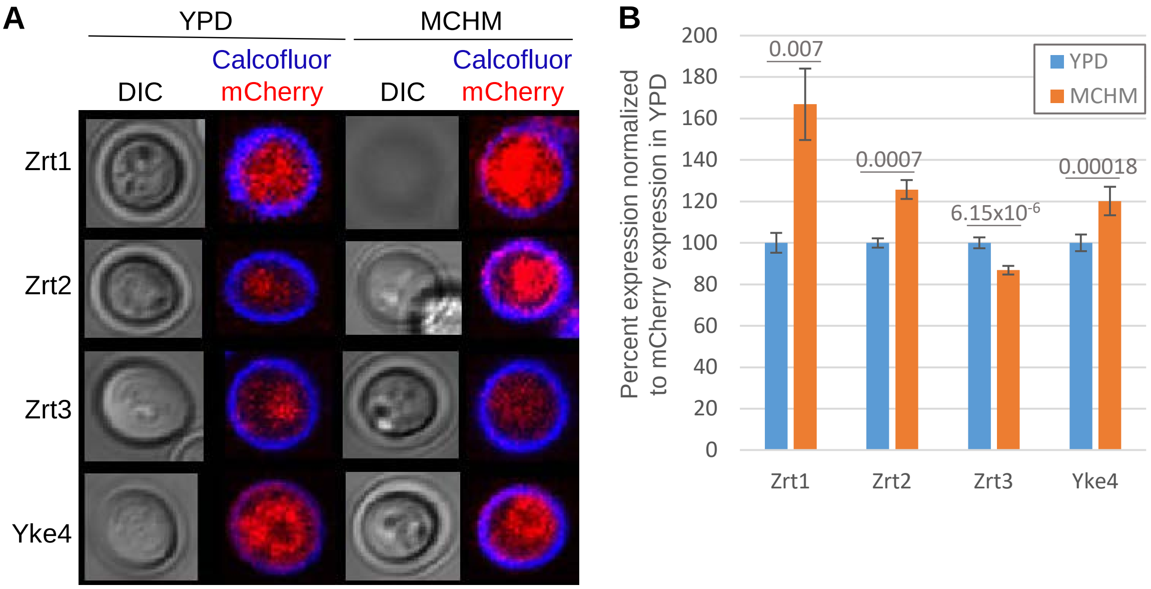


**Supplemental Figure 6 Pupo 2018**
